# Evidence for a role for BK channels in the regulation of ADAM17 activity

**DOI:** 10.1101/811000

**Authors:** Minae Yoshida, Dean Willis

## Abstract

Large-conductance voltage and calcium activated channels, K_Ca_1.1, have a large single conductance (~p250) and are highly selective for potassium ions. As a result they have been termed big potassium channels (BK channels). Because of the channel’s ability to integrate multiple physical and chemical signals they have received much attention in excitable cells. In comparison they have received relatively little attention in non-excitable cells in those of the immune system. Here we report evidence that the BK channel regulates ADAM17 activity. Upon macrophage activation, BK channels translocate to the cell membrane. Genetic or pharmacological inhibition of the cell membrane BK channels resulted in elevated TNF-α release and increased metalloproteinase a disintegrin and metalloproteinase domain 17 (ADAM17) activity. Inhibitors of BK channels also increased IL-6Rα release, a second ADAM17 substrate. In comparison, a BK channel opener decreases TNF-α release. Taken together, our results demonstrate a novel mechanism by which ion channel regulates ADAM17 activity. Given the broad range of ADAM17 substrates, this finding has implications in many fields of cell biology including immunology, neurology and cancer biology.

## Introduction

Macrophages are an essential component of the immune system *(1, 2)* with activated macrophages being a major source of the cytokines *(3)*. Due to the importance of cytokines in health and disease the signaling cascades which contribute to their expression and release by macrophages have been extensively researched. TNF-α is a potent pro-inflammatory cytokine released from macrophages. The importance of this cytokine is demonstrated by the clinical efficacy of anti-TNF-α drugs in a variety of inflammatory diseases such as rheumatoid arthritis. One aspect of TNF-α biology which as recently become apparent is that released soluble TNF-α, and non-released, membrane bound TNF-α (mTNF-α), contribute to different cellular signaling cascades via different receptors or reverse signaling*(4–7)*. This adds an additional layer of complexity to TNF-α biology and highlights the importance of mechanisms by which cytokines and their receptors are released from the cell membrane

Sheddases are membrane bound enzymes which cleave membrane bound proteins allowing there release. ADAM17 is a member of this family of proteases, which mediates the release of a diverse range of proteins including cytokines, growth factors, receptors and adhesion molecules from the cell surface *(8)*. The release of these molecules by ADAM17 is implicated in intercellular communication pathways in many fields of biology including immunology, neurology, embryology and tumor growth *(8–10)*. TNF-α is one of the best known examples of a molecule whose release is regulated by ADAM17. Therefore the mechanisms by which ADAM17 activity is regulated would be of interest.

Given the wide expression of ion channels in macrophages *(11, 12)* and their intimate involvement in membrane biology we speculated that ion channels may have a role in regulating cytokine release.

The large conductance voltage-and calcium-activated potassium channel (BK channel) is characterized by a large single channel conductance and consists of 4 core channel-forming α-subunits (BKα). This channel opens in response to a rise in intracellular Ca^2+^ and/or membrane voltage *(13)*. However, various combinations of auxiliary subunits, either β1-4 and/or γ1-4 modify the channel’s opening properties *(13–15)*.

In this paper, we present evidence that BK channels are upregulated on the cell membrane of macrophages and regulate the release of TNF-α from the cell membrane with evidence pointing to role for ADAM17 modulation. This paper highlights a new mechanism by which macrophages regulate the release of TNF-α and due to the cellular importance of other ADAM17 substrates may be of interest to researchers in a variety of fields.

## Results

### Silencing BK channels increases TNF-α release and ADAM17 activity

RAW264.7 cells are a standard mouse macrophage cell line in which TLR4 cell signaling events and TNF-α production are well characterized *(16)*. Western blot analysis of whole cell lysates from resting RAW264.7 macrophages detected the presence of the essential BKα subunit at 120 kDa (Fig. 1A). To evaluate the role of BK channels in macrophages, we employed siRNA to knock-down the expression of BKα. Macrophages are notoriously difficult to transfect *(17)*. In attempt to optimize transfection efficiency while minimizing toxicity and cell activation, we investigated a range of transfection reagents, siRNA sequences, concentrations and incubation times in RAW264.7 macrophages before adopting the protocol described in Material and Method. Even after the optimization experiments, the overall silencing effect on whole cell BKα in RAW264.7 macrophages was limited to a 25% reduction in protein expression compared to non-silencing siRNA controls (Fig. 1A). However, it was found that BKα silencing siRNAs had a greater effect on the expression of plasma membrane associated BKα, approximately 65% reduction compared to controls (Fig. 1B). These results suggest that the siRNA protocol adopted for this study preferentially affected the cell surface expression of BK channels compared to global BKα expression. Bearing this observation in mind, we next examined the effect of BKα silencing on LPS induced macrophage TNF-α production.

**Fig. 1.**
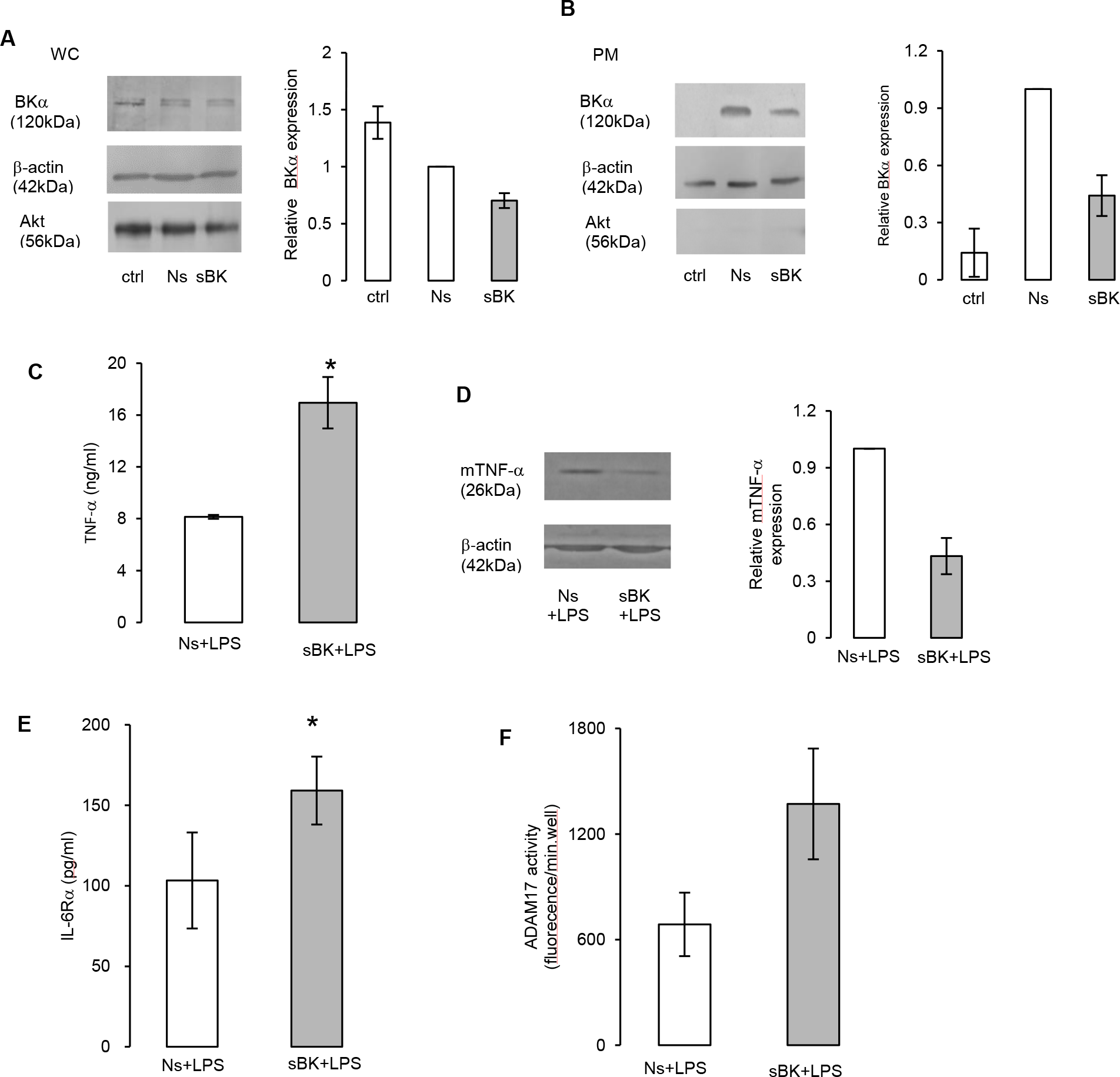
BK channel silencing increased ADAM17 activity and enhanced secretion of TNF-α and IL-6Rα.. (**A**)&(**B**) Western blot analysis of whole cell and plasma membrane proteins from siRNA treated macrophages. The cells were non-treated (ctrl), received non-silencing siRNA (Ns) or silencing siRNA for BKα genes (sBK) for 72 hours. BKα silencing effect in (**A**) whole cell (WC) and (**B**) plasma membrane (PM) lysate. Densitometry analysis of BKα expression in BKα silenced cells relative to non-silencing siRNA. Mean+/−SEM (n=3). (**C**)-(**F**)After treatment with siRNAs, macrophages were activated with 150 ng/ml LPS for 4 hours. (**C**) TNF-α and (**E**) IL-6Rα release was analyzed by ELISA. Mean+/−SEM (n=4). (**D**) Western blot analysis of mTNF-α expression. Densitometry of mTNF-α expression in BKα silenced cells relative to non-silencing siRNA. Mean+/−SEM (n=3). (**F**) ADAM17 activity in BKα silenced (sBK) macrophages. Mean+/−SEM (n=3). *p<0.05.

Initial experiments established that 4 hour stimulation with 150 ng/ml LPS gave the maximal levels of TNF-α release from RAW264.7 cells. Silencing of BK channels increased TNF-α secretion from these cells by 107% when compared to non-silencing controls (Fig. 1C). TNF-α is initially expressed as a membrane bound 26 kDa protein, mTNF-α. ADAM17 cleaves mTNF-α at its defined ecoto-domain amino-acid sequence to release the soluble 17 kDa TNF-α *(18–20)*. BKα silencing resulted in a 55% decrease in mTNF-α as determined by Western blot analysis (Fig. 1D). The release of IL-6Rα, another substrate for ADAM17 *(8)*, was also elevated after BKα silencing (Fig. 1E). We therefore hypothesized that the BK channel regulates ADAM17 activity. To assess this directly, we used a fluorogenic substrate for the ADAM17. ADAM17 activity during LPS activation was almost doubled in BK silenced macrophages compared to non-silencing controls (Fig. 1F). These results support the hypothesis that BK channels on the plasma membrane can limit the activity of ADAM17.

### LPS treatment upregulates BK channel expression on the plasma membrane

To provide further evidence for the siRNA data demonstrating a role for BK channel in regulating ADAM17 activity, we next used a pharmacological approach. BK channels are found in a variety of cellular locations. These include the plasma, mitochondrial and nuclear membranes, where they have differing cellular functions *(13, 21–24)*. Because the siRNA silencing experiment appeared to preferentially inhibit BKα expression on the plasma membrane and this location would be a logical site of the channels to regulate ADAM17 activity, we hypothesized that it is plasma membrane located BK channels that regulate ADAM17 activity and not intracellular channels. To selectively block the plasma membrane located channels, we utilized a membrane impermeable BK channel inhibitor, iberiotoxin (IbTX). IbTX is a 37 amino acid peptide, which blocks the BK channel from the extracellular side and is highly selective at concentrations below 100 nM *(25)*. Using IbTX allowed us to selectively inhibit plasma membrane BK channels by pharmacological means.

IbTX had no significant effect on TNF-α secretion or ADAM17 activity in resting macrophages or in macrophages stimulated for 4 hours with 150 ng/ml LPS (Fig. 2, A and B). Importantly this lack of IbTX effect was correlated with a low level of BK channel expression on the plasma membrane of resting/non-treated macrophages (Fig. 1B & Fig. 2C). Taken together, the data suggested that under resting/non-treated conditions, macrophages do not express many BK channels on their plasma membrane or do not participate in early TNF-α release. The presence of several types of K^+^ channels has been shown in human and rodent macrophages and macrophage cell lines. These K^+^ channels include K_V1.5_, K_V1.3_, K_Ca3.1_, K_ir2.1_ and the BK channel with the differentiation/activation of macrophages being reported to increase the expression of K^+^ channels on their cell surface *(11, 12, 26, 27)*. Indeed, even though we attempted to minimize RAW264.7 activation in our siRNA protocol, our data suggested that treatment with the adopted protocol increased plasma membrane BK channel expression in RAW264.7 macrophages (Fig. 1B). This observation again suggested that BK channel cellular expression changes after the macrophage activation. Therefore we decided to test whether LPS activation affected the channel expression and cellular location. As we found that highly activated macrophages are particularly difficult to collect electrophysiological recordings from, we stimulated cells with a lower, submaximal, dose of LPS, 10 ng/ml for up to 24 hours.

**Fig. 2.**
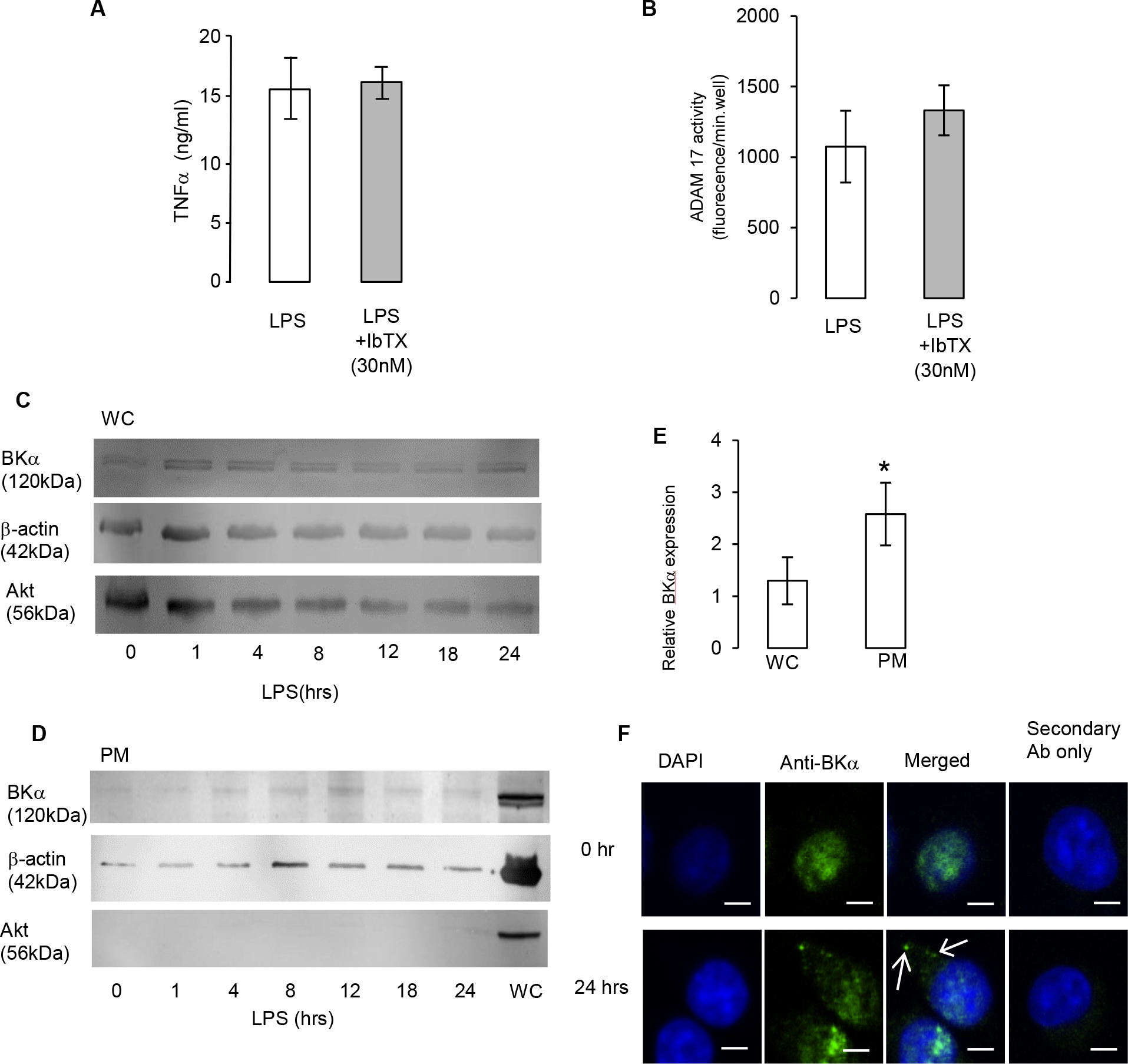
Macrophages express plasma membrane BK channels after LPS activation. Plasma membrane BK channels expression appear after LPS treatment. (**A**) TNF-α release measured by ELISA and (**B**) ADAM17 activity in macrophages received 150 ng/ml LPS for 4 hours with/without IbTX. Mean+/−SEM (n=3). Western blot analysis of BKα expression in (**C**) whole cell (WC) and (**D**) plasma membrane (PM) lysate from macrophages treated with 10 ng/ml LPS for up to 24 hours. (**E**) Densitometry of BKα expression after 24 hours of LPS treatment relative to 0 hours. Mean+/− SEM (n=3). (**F**) Immunolocalization of BKα. Staining for DAPI, anti-BKα antibody (green), merged image of DAPI and anti-BKα staining and secondary antibody only control with DAPI. Scale bar, 5μm. Arrows indicate BKα staining on the plasma membrane. *p<0.05.

At the whole cell level, the expression of BKα protein remained relatively constant for 24 hours of 10 ng/ml LPS treatment (Fig. 2C). Immunofluorescence imaging of resting macrophages indicated that the plasma membranes appeared relatively free of the BKα protein, instead BKα positive staining was predominantly associated with the cytoplasm and nucleus of these cells (Fig. 2F). This cellular distribution of BKα was confirmed by Western blot analysis (Fig. 2D). A previous study has demonstrated the expression of nuclear BK channels in neurons and proposed a role for the nuclear BK channel in controlling CREB-mediated transcription *(22)*. This observation of nuclear BK channel in macrophages has not been previously reported and it will be of interest to investigate its role in macrophage biology in the future. To further evaluate the expression of functional BK channels on the plasma membrane, we performed whole cell voltage clamp experiments of resting macrophages. Initial traces using a non-selective potassium channel blocker, TEA, demonstrated the sparsity of K^+^ channels on plasma membranes (Fig. 3A). Furthermore, only 8.7% of resting macrophages exhibited IbTX-sensitive K^+^ current (Fig. 3B). These results indicated that the majority of resting macrophages do not express BK channels on their cell surface.

**Fig. 3.**
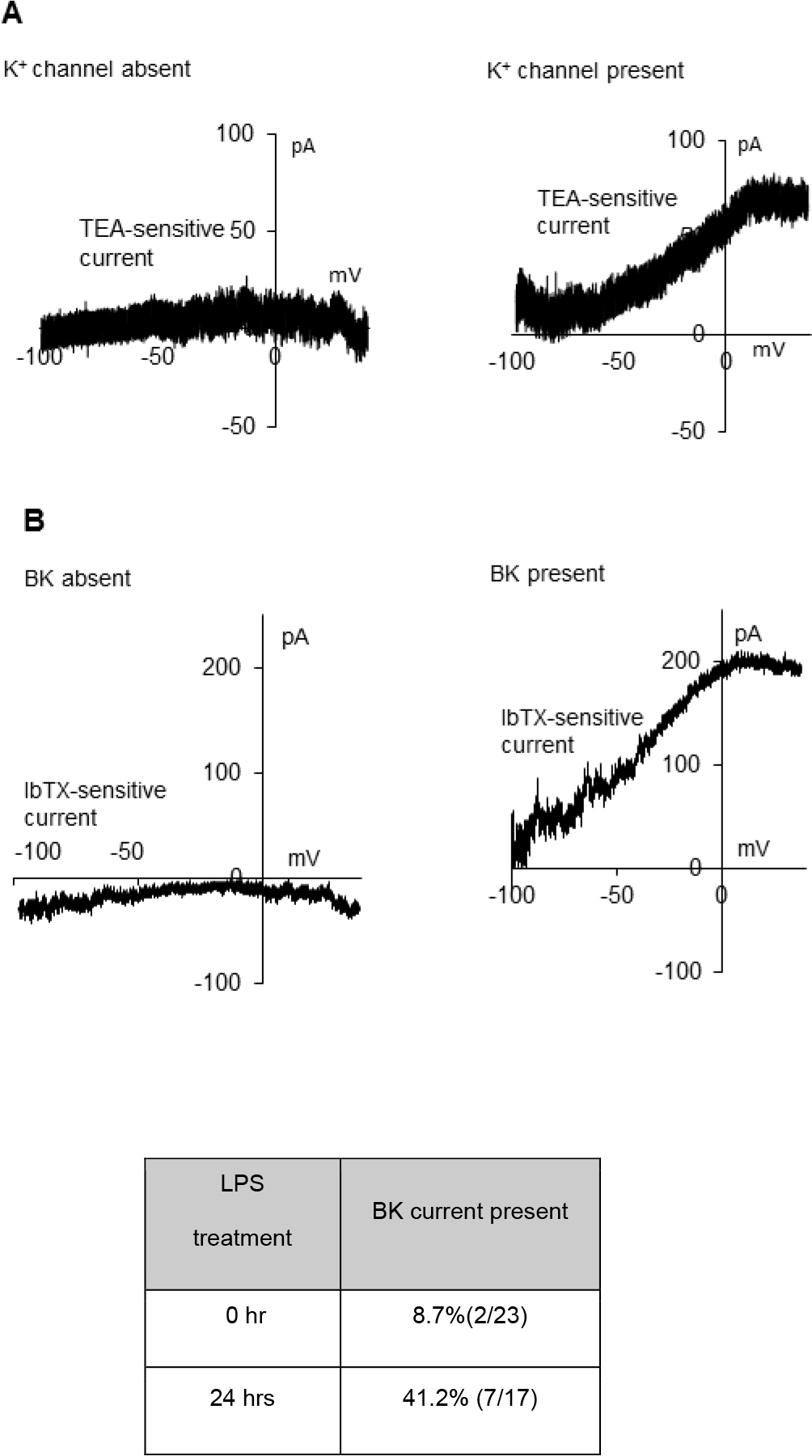
BK channel activity appears on the cell surface of macrophages after LPS activation. Representative traces of whole cell recordings from resting or 24 hours 10 ng/ml LPS stimulated macrophages. Voltage ramp (−100 mV to +40 mV, duration 10 s) were applied in presence or absence of 3 mM TEA or 30 nM IbTX. Representative graphs of (**A**) TEA- and (**B**) IbTX-sensitive current. These drugs were applied in the extracellular solution. The ramp protocol was run ≥6 times with each drug treatment and these traces were averaged. Sensitive currents were obtained by subtraction of traces with drug treatment from control traces and plotted against membrane voltage.

While total cell BKα protein stayed at similar levels, the plasma membrane expression of BK channel increased during the 24 hours of 10 ng/ml LPS treatment (Fig. 2 D and E). After 24 hours of LPS stimulation, BKα, expression in the plasma membrane had increased by 150% compared to 0 hours whereas whole cell BKα expression only increased by 30% (Fig.2C). Immunofluorescence imaging also confirmed the presence of BKα expression on the plasma membrane between 18-24 hours of LPS treatment (Fig. 2F) with a staining pattern indicative of channel clustering *(28)*. 24 hours after LPS stimulation, whole cell recordings showed the presence of TEA-sensitive K^+^ channels (Fig. 3A), and IbTX blocked K^+^ currents in 41.2% of these macrophages (Fig. 3B). This demonstrates that LPS causes 5 fold-increase in the number of cells expressing functional BK channels on their plasma membrane.

### Pharmacological modulation of BK channel activity regulates TNF-α release and ADAM17 activity

To further study the BK channel on the plasma membrane and its potential effect on ADAM17, we treated macrophages with 10 ng/ml LPS for 24 hours. A higher proportion of these macrophages express plasma membrane BK channels than non-stimulated, resting macrophages. These LPS treated macrophages were termed ‘conditioned macrophages’. These cells were further stimulated with 150 ng/ml LPS for 4 hours in absence/presence of 30 nM IbTX to investigate the function of plasma membrane BK channels on ADAM17 substrates (Fig. 4A).

**Fig. 4.**
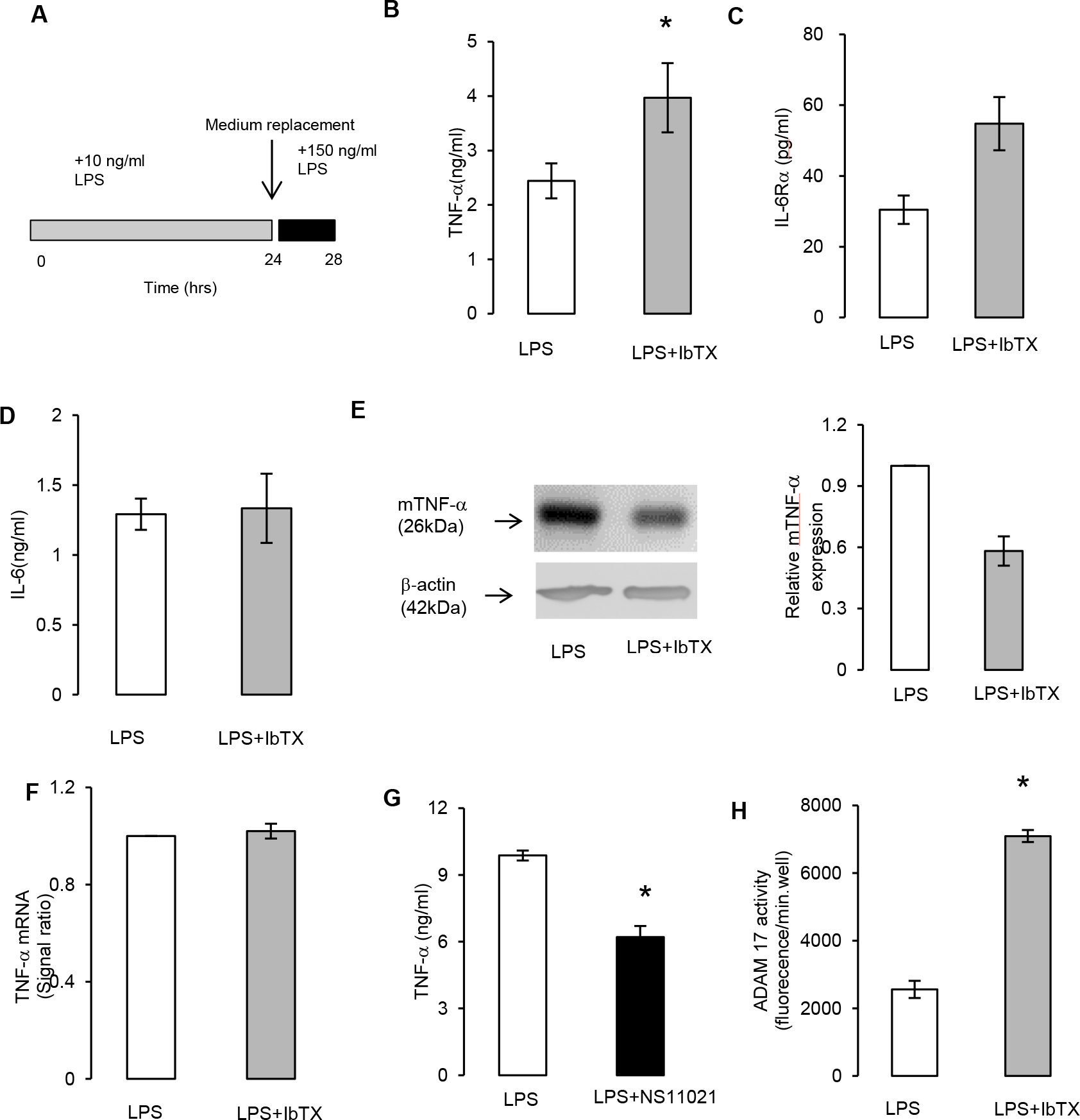
Pharmacological inhibition of BK channels increased ADAM17 activity and the release of TNF-α, and IL-6Rα. (**A**) Conditioned macrophages were prepared by stimulating cells with 10 ng/ml LPS for 24 hours. Subsequently the media was replaced and the conditioned cells were activated with 150 ng/ml LPS for 4 hours with/without 30 nM IbTX. (**B**) TNF-α, (**C**) IL-6Rα and (**D**) IL-6 release in were analyzed by ELISA. Mean+/−SEM (n=3). (**E**) Western blot analysis of mTNF-α expression and densitometry of mTNF-α expression relative to LPS only control group. (**F**) TNF-α mRNA was quantified by RT-qPCR. Mean+/−SEM (n=3). (**H**) ADAM17 activity. Mean+/−SEM (n=3). (G) Effect of 100 nM NS11021 on TNF-α release. *p<0.05.

IbTX treatment resulted in a 55% increase in TNF-α, release from LPS stimulated conditioned macrophages (Fig. 4B) and corresponded to a 44% decrease in mTNF-α compared to controls (Fig. 4E). RT-qPCR showed no change in TNF-α mRNA expression after IbTX treatment (Fig. 4D). This data supports a post-transcriptional mechanism by which BK channels control TNF-α release in these cells. Furthermore, application of a BK channel opener, 100 nM NS11021, significantly decreased TNF-α release by 37% (Fig. 4G). IbTX increased the release of a second substrate of ADAM17, IL-6Rα (Fig. 4C), but did not influence the release of IL-6, which is not a substrate of ADAM17 (Fig. 4D). Finally IbTX treatment increased ADAM17 activity by 150% (Fig. 4H). Taken together, these results support a role for plasma membrane BK channels in regulating ADAM17 activity.

## Discussion

Ion channels are widely expressed in macrophages, however their role in these cells remains to be fully elucidated *(11)*. While investigating the function of ion channels within a standard macrophage cell line, we identified a role for BK channels in regulating the release of TNF-α from these cells. Our work suggests that after stimulation, macrophages upregulate the expression of functional BK channels on their plasma membrane and once present, they participate in the regulation of ADAM17 activity. These results highlight a new mechanism by which BK channels can regulate the release of proteins from cell membranes.

The activation/differentiation of immune cells has previously been shown to upregulate the expression of K^+^ channels on the plasma membrane *(11, 26)*. In line with this general observation, our study showed that macrophage activation led to an increase in BK channel expression on the plasma membrane (Fig. 2D). It is known that the BK channel participates in different cellular processes depending on its cellular location. The location of BK channels is therefore an important factor when investigating the function of these channels in whole cells and in vivo. For example, data obtained from selective inhibition of BK channels on the plasma membrane may be different from pan BKα suppression as seen in knockout animals *(29)*. In our study, plasma membrane BK channels appeared to be inhibited preferentially by siRNA (Fig. 1 A and B) or specifically inhibited by the membrane impermeable channel blocker IbTX.

The major finding of this study is that the plasma membrane BK channel regulates ADAM17 activity. ADAM17 is implicated in the release of a broad range of proteins from cell membranes. It has been reported that there are at least 76 protein substrates for this enzyme, which are involved in multiple cell/tissue functions, however whether these are all unique substrates of ADAM17 requires further variation *(8)*. A central question is what mechanism(s) cells employ to allow the precise temporal and spatial release of a diverse set of ADAM17 substrates. Mechanisms believed to contribute to the control of ADAM17 activity include; cleavage of the enzyme pro-domain by furin proteases *(30)*, translocation of ADAM17 to plasma membrane by iRhom2 *(31–33)* and the conversion of ADAM17 dimers to active monomers by dissociation of the endogenous inhibitor TIMP-3 *(34)*. Recently the externalization of phosphatidylserine has been shown to facilitate the interaction between the active site of the ADAM17 and cleavage site of its substrates via negatively charged residues on the phospholipid *(35)*. In this paper we did not address the mechanisms by which BK channel regulated ADAM17 activity. However this could feasibly by multiple levels. Importantly the co-localization of BK channels with ADAM17 and its substrate on the plasma membrane could represent a precise control mechanism by which proteins are released in a temporal and spatial fashion from the macrophage.

What are the biological consequences of the BK channel regulating the release of TNF-α and IL-6Rα from macrophages? Macrophages are a major producer of TNF-α and play a central role in inflammatory reactions *(3, 8)*. TNF-α signals via two receptors TNFR1 and TNFR2, importantly, soluble TNF-α and mTNF-α signals via TNFR1, while only mTNF-α signals via TNFR2 *(4–6)*. TNF-α is considered to be a potent pro-inflammatory cytokine and TNF-α inhibitors are used to treat a variety of inflammatory conditions including rheumatoid arthritis, ankylosing spondylitis, inflammatory bowel disease and psoriasis *(5, 6)*. While these drugs have been successful, there is increasing evidence in tissue specific models of inflammation which suggests that mTNF-α signaling via TNFR2 is anti-inflammatory/tissue protective. Evidence for this tissue protective role of TNF-α in humans comes from the observations that TNF-α inhibitors exacerbate Multiple sclerosis and that some patients with Rheumatoid arthritis who have taken these TNF-α inhibitors have developed Multiple sclerosis like pathology *(36–38)*. It has therefore been suggested that inhibiting soluble TNF-α signaling while maintaining mTNF-α signaling and/or limiting the cellular source of soluble TNF-α may be a superior therapy to pan TNF-α inhibition *(39)*. This paper provides evidence that the opening of BK channels can inhibit the conversion of mTNF-α to soluble TNF-α in macrophages; and from a potential therapeutic point of view, we demonstrate that a BK channel opener can inhibit TNF-α release (Fig. 4F). BK channels also regulate the release of IL-6Rα. Membrane bound IL-6Rα participates in classical signaling with IL-6, which is considered to be essential for regenerative and anti-bacterial actions of this cytokine; whereas soluble IL-6Rα participates in trans-signaling with IL-6, which contributes the pro-inflammatory/deleterious effects of the cytokine *(40)*. Our results would suggest that BK channel openers could favor the expression of the membrane-bound forms of mediators, e.g. mTNF-α and IL-6Rα, and shift the balance away from the soluble form of this proteins. This would result in a tissue environment which was anti-inflammatory/regenerative in nature. Interestingly some endogenous openers of BK channels, including carbon monoxide and epoxyeicosatrienoic acid *(41, 42)*, have been shown to have anti-inflammatory actions and to participate in the resolution of inflammation *(43)*. Additionally, we have demonstrated that BK channels are upregulated on the plasma membrane after LPS activation. This phenomenon could allow the macrophage to alter their phenotype between pro-inflammatory and anti-inflammatory states. It is therefore possible that BK channels expression contributes to the adaptation of macrophages in different tissue/micro environments.

In conclusion BK channels and ADAM17 are widely expressed, with the enzyme regulating the release of diverse set of substrates. This study identities a novel interaction between the channel and the enzyme which may open up new avenues of research and possible therapeutic opportunities.

## Materials and Methods

### Reagents

Ultrapure LPS (Invivogen), tetraethylammonium (TEA) (Sigma) and Iberiotoxin (IbTX) (Alomone Labs) were reconstituted in sterile distilled water. NS11021 (Sigma), GM6001 (Enzo Life Science) and TAPI-0 (Enzo Life Science) were reconstituted in DMSO. Where appropriate, DMSO was added to control groups in case of NS11021, the final concentration of 0.015% and in the ADAM17 activity assays 0.1%. Other chemicals are purchased from Sigma unless otherwise stated.

### Antibodies and staining reagents

Anti-mouse BKα mouse monoclonal antibody (L6/60, Millipore; 1 in 1000 Wetsern blot (WB), 1 in 750 in Immunofluorecence (IF)), anti-ADAM17 rabbit polyclonal antibody (ab2051, Abcam; 1 in 1000 WB), anti-Akt rabbit polyclonal antibody (Cell Signaling Technology; 1 in 1200 WB), anti-mouse TNF-α rabbit monoclonal antibody (D2D4, Cell Signaling Technology; 1 in 1000 WB) and anti-β-actin rabbit polyclonal antibody (D6X8, Cell Signaling Technology; 1 in 1500 WB) were used. Goat anti-rabbit HRP (Cell Signaling Technology; 1 in 4000 WB) and rabbit anti-mouse IgG H&L (ab97046, Abcam; 1 in 5000 WB) were used as secondary HRP conjugated antibodies in Western blotting. F(ab)’ fragment affinity-purified unconjugated goat anti-mouse IgG (Thermo Fisher Scientific; 1.25 μg/ml) and goat anti-mouse IgG (H&L) Alex FluorR 488 conjugate (Thermo Fisher Scientific; 1 in 2000 IF) were used in immunofluorescence imaging.

### Cell culture

RAW264.7 murine macrophages were obtained from the European Collection of Cell Culture. The cells were cultured in Dulbecco’s Modified Eagle Medium (DMEM) (Thermo Fisher Scientific) containing 10% fetal bovine serum, 100 unit/ml penicillin and 100 μg/ml streptomycin in humidified incubator at 37°C with 5% CO_2_. The cells were passaged when 80% confluent. Mycoplasma contamination was routinely tested for by DAPI staining or using a mycoplasma detection kit, Mycoprobe (R%D Systems) following the manufacturer’s instruction. MTT tests confirmed that the treatments did not cause cell death.

### Plasma membrane isolation

Plasma membrane protein was isolated using Pierce Cell Surface Protein Isolation Kit (Thermo Fisher Scientific) according to manufacturer’s protocol. Each sample was prepared using 3×10^7^ cells. After the surface protein was labelled with 1 mg/ml Sulfo-NHS-SS-Biotin (Thermo Fisher Scientific), the cells were lysed in PBS containing protease inhibitor cocktail (consisting of AEBSF, aptrotinin, bestatin hydrochloride, E64,EDTA and leupeptin hemisulfate salt) and 1% IGAL-CA60 by sonication at 4°C. The labelled proteins in post nuclear supernatants were captured with Neutroavidn agarose. Captured proteins were eluted with SDS-PAGE sample buffer, containing 50 mM DTT. The protein contents in sample elute were equilibrated to each other after protein measurement with a Bradford assay. The absence of the cytosolic protein, Akt, in Western blots was used to validate plasma membrane isolation process.

### Western blot analysis

Whole cell and nuclei isolate from RAW264.7 macrophages were lysed by sonication in PBS containing protease inhibitor cocktail at 4°C. Protein contents in the lysate were measured by Bradford assay and protein equivalents loaded in SDS-PAGE sample buffer. The samples were separated on SDS-PAGE, and transferred to nitrocellulose membranes (Thermo Fisher Scientific). Membranes were blocked with 5% non-fat milk and 0.05% BSA in Tris-buffered saline (TBS) with 0.1% Tween-20 for 2 hours, and stained with primary antibodies for BKα, TNF-α, ADAM17, Akt or β-actin overnight at 4°C. Membranes were incubated with horseradish-conjugated anti-IgG antibodies. Immunoreactive bands were detected using chemiluminescence with Pierce ECL2 Western Blot Substrate (Thermo Fisher Scientific) and scanned by a phosphor-imager (Typhoon 9410 variable made imager, GE Healthcare). Experiments were repeated at least 3 times, representative blots are shown in figures. Densitometry analysis was performed using Image Quant 5.2 (Molecular Dynamics). The signal of the target protein was normalized to β-actin. Signal ratio to the relevant control group is expressed as mean+/−SEM (n=3). n: number of blots used for the densitometry analysis.

### Electrophysiology

Whole cell voltage clamp recordings were performed at room temperature. Macrophages were plated on a glass recording chamber. Pipettes were made using thin walled borosilicate glass (Harvard), heat-pulled and polished. When filled with pipette solution, the pipette had a resistance between 3-4 MΩ. The pipette solution was composed of 120 mM K-acetate, 20 mM HEPES, 10 mM KCl, 4 mM CaCl_2_, 1 mM EGTA, 1 mM MgCl_2_ and had 1.6 μM free Ca^2+^, pH=7.4 and osmolality between 250-300 mOsmol. The extracellular solution was Kreb’s: 125 mM NaCl, 26 mM NaHCO_3_, 2.5 mM KCl, 1.26 mM NaH_2_PO_4_ H_2_O, 25 mM D-glucose, 1 mM CaCl_2_. Currents were recorded using an Axopatch 200B amplifier (Axon Instruments) at the sample rate of 5 kHz and filtered at 1 kHz with Digitata 1332A (Molecular Devices). Once a high resistance seal was formed, a ramp protocol (25 s with 10 s sweep ranging from −40 to 100 mV) was applied. 3 mM TEA and 30 nM IbTX were applied in extracellular solution. pH of extracellular solutions were adjusted to 7.4. TEA- and IbTX-sensitive currents were obtained by subtraction, and representative traces were plotted against membrane voltage. Recordings were carried out at room temperature. pCLAMP software (Molecular Devices) was used to monitor current magnitude and duration, establish voltage ramp protocols and analyze the data.

### Cytokine assays

RAW264.7 macrophages were plated at 2.5×10^5^ cells per well on 24 well plates. The cells were incubated overnight. The following day, the media was replaced and the cells received various drug treatments. TNF-α and IL-6Rα in media duplicates were quantified using ELISA kits (R & D Systems) according to manufacturer’s protocol. The experiments were repeated at least 3 times with n=3 or 4. Graphs show mean+/−SEM of representative data and are expressed as mean+/−SEM (n=3, 4). n: number of biological replicates per treatment group in each experiment.

### RNA extraction and cDNA synthesis

Total RNA was extracted using RNAsay mini-kit (Qiagen). The quality and purity of RNA were assessed by OD260/280 ratio, measured using a spectrophotometer (DU800, Beckman Coulter). Complementary DNA (cDNA) was synthesized using the Superscript III Reverse transcriptase kit (Thermo Fisher Scientific) with random primers (Invivogen). 2 μg of total RNA were used per construct.

### Real time-quantitative PCR

RT-QPCR was performed on cDNA using the Taqman custom gene expression assay (Thermo Fisher Scientific) and the ABI Prism 7900HT sequence detection system (Thermo Fisher Scientific). The probes for mouse TNF-α (Mm00443260_g1) and mouse β-actin (Mm00607939_s1) (Thermo Fisher Scientific) were used. The assay was set up in accordance with Taqman protocols and TNF-α expression level was normalized to β-actin. Experiments were repeated 3 times with n=3. n: number of biological replicates per treatment group in each experiment. Graph shows representative data and is expressed as mean+/−SEM (n=3).

### ADAM17 activity assay

Proteolytic activity of ADAM17 was assessed by monitoring cleavage of TACE substrate II (fluorogenic) from live cells by adapting the method described by Alvarez-Iglesias et al *(44)*. RAW264.7 macrophages were plated at 4.8×10^4^ cells per well on a 96 well fluorescence cell culture plates. Assay was carried out with phenol red free DMEM supplemented with 4 mM L-glutamine (Thermo Fisher Scientific). Three hour before the assay, the macrophages received combinations of 150 ng/ml LPS and 30 nM IbTX. TACE substrate II fluorogenic (BML-P228, Enzo Life Sciences) was reconstituted in DMSO, directly dissolved in media and applied to the cells at the final concentration of 5 μM. Fluorescence was measured for 60 minutes using a fluorescence plate reader (Mithras LB 940, Thermo Fisher Scientific). Samples containing no TACE substrate acted as negative control. A general inhibitor for membrane metalloproteases, GM6001, and a selective inhibitor for ADAM17, TAPI-0, were used to validate that the fluorescence signal was due to the ADAM17 activity. Data shows per minute increase of fluorescence signal as mean+/−SEM (n=3, 4). n: number of biological replicates per treatment group in each experiment. Assays were repeated at least 3 times with n=3 or 4 and graphs show representative data.

### Gene silencing

BKα gene silencing was carried out using Lipofectamine® RNAimax reagent (Thermo Fisher Scientific) and two silencing siRNA for mouse BKα (Target sequences CAC AAT GTC TAC AGT GGG TTA and CAG TTT CTG AAT ATC CTT AAA) from Qiagen. The two siRNA were mixed in equal ratio. Negative control groups received non-silencing siRNA (Target sequence AAT TCT CCG AAC GTG TCA CGT) from Qiagen. The cells were incubated with the siRNA for 72 hours. T25cm^2^ cell culture flasks were used for transfection, and each flask had 1.25×10^6^ cells. 90 pmol of total siRNA and 27 μl of Lipofectamine®6 RNAiMAX were used per flasks containing 6 ml of media.

### Immune fluorescence staining

For immunolocalization of BKα, macrophages were plated at 1.3×10^4^ cells per well onto glass circular coverslips and treated with 10 ng/ml LPS for up to 24 hours. The cells were fixed with 4% paraformaldehyde. The coverslips were blocked with 1.25 μg/ml F(ab)’ fragment affinity-purified unconjugated goat anti-mouse IgG, 10% goat serum and 4% bovine serum albumin (BSA) in PBS for 1 hour followed by 0.01% Triton X-100 for 5 minutes. The coverslips were then incubated with anti-BKα antibody diluted in 5% goat serum and 2% BSA in PBS at 4°C overnight, followed by secondary antibody, goat anti-mouse IgG (H&L) Alex FluorR 488 conjugate for 2 hours, and DAPI. Coverslips were mounted on glass slides with Vectashield (Vector Laboratories). Fluorescence was visualized using a confocal microscopy (Zeiss LSM510 Meta, Carl Zeiss). Experiments were repeated at least 3 times and figures show representative images.

### Statistical analysis

Prism (GraphPad Software) was used for statistical analysis. Statistical significance was determined by Student t-test for the comparison between two groups and for the comparison among three or more groups, significance was tested by one-way ANOVA, followed by Bonferroni t-test. Either case, a P-value of less than 0.05 was considered to be significant.

